# Comprehensive circular RNA expression profiles in intermuscular fat from lean and obese pigs

**DOI:** 10.1101/546390

**Authors:** Zhiguo Miao, Jinzhou Zhang, Shan Wang, Paneng Wei

**Affiliations:** College of Animal Science and Veterinary Medicine, Henan Institute of Science and Technology, 453003, Xinxiang, Henan, China

**Keywords:** Pigs, CircRNA, Intermuscular fat, Expression profiles

## Abstract

Circular RNA (circRNA) plays an important regulatory role in development and differentiation. Intermuscular fat in pork affects the tenderness and juiciness of the meat. In this study, we investigated the performances of Landrace (lean) and Jinhua (obese) pigs at the fattening period and explored the expression profile of circRNAs in intermuscular fat from the two breeds by Illumina high-throughput sequencing. we identified 5 548 circRNAs, specifically 2 651 (47.78%) in the Jinhua pigs, and 2 897 (52.22%) in the Landrace pigs. A totale of 809 differentially expressed circRNAs were observed between the the Jinhua and Landrace pigs, but only 29 of these circRNAs showed significant difference (19 upregulated and 10 downregulated). All 1 306 unigenes and 27 differentially expressed unigenesinvolved in lipid transport and metabolism; replication, recombination and repair; and signaling pathway were annotated. A total of 550 target miRNAs were perfect seed matches and 20 522 target genes were foundBy RNAhybrid and miRanda software prediction. Results from real-time quantitative PCR also confirmed the differential expression of 13 mRNAs between the two pig breeds. This study provides comprehensive expression profiles of circRNAs in *Sus scrofa* adipose metabolism and development, which can be used to clarify their functions.

**Summary statement:** The paper explored the expression profile of circRNAs in intermuscular fat from Landrace and Jinhua pigs at the fattening period, to provide insights into circRNAregulation in animal adipose metabolism.

## 1. Introduction

Circular RNA (circRNA) plays a novel central regulatory role in RNA metabolism (Cortés-López M and Miura, 2016). CircRNA is a noncoding RNA that forms covalently closed continuous loops, engendering from the backsplicing of exons, introns, or both to form exonic or intronic RNA (Cortés-López and Miura, 2016). CircRNA is predominantly produced by back-splicing reactions that covalently link the 3 ′end of an exon to the 5′ end of an upstream exon (Vicens and Westhof, 2014). Although identified over 30 years ago, circRNA was thought to be an error in RNA splicing. Recent studies have confirmed that circRNA acts as a transcription regulator and that it has other functions (Memczak *et al.*, 2013; Hansen *et al.*, 2013; Venø *et al.*, 2015).

CircRNA is a new regulator of gene expression. Many studies focused on the spatiotemporal regulation of circRNA in animal (Venø *et al.*, 2015; Zeng *et al.*, 2017). CircRNA yields specifically expressed patterns in different tissues, cell types, or developmental stages, showing its important roles in developmental regulation (Venø *et al.*, 2015; Salzman, 2016). Hansen *et al.* (2013) found that circRNAs can regulate the expression of corresponding genes at the post-transcription level via the microRNA (miRNA) pathway. Cerebellar degeneration-related protein 1 transcript (**CDR1a**), which maybe the best-characterized mammalian circRNA, is highly abundant in neurons, circularizing into circRNA harbored 73 conserved binding sites for targeting miR-7(Jens, 2013). Furthermore, it interacts with miR-7 and strongly suppresses miR-7 activity, which allows mRNAs to escape degradation following miRNA binding and then increases the levels of miR-7 targets. Ashwal-Fluss *et al.* (2014) observed that the biogenesis of MBL1 circRNA derived from the muscle-blind locus in *Drosophila* is stimulated by the Mbl1 protein and that MBL1 circRNA interacts with Mbl1, thereby reducing *MBL1* mRNA production and controlling Mbl1 expression. However, the specific mechanism underlying circRNA biogenesis remains unclear.

Across its domestication, pig (*Sus scrofa*) is an important domesticated farm animal that provides protein for humans. The domestic pig is increasingly being used as a nonrodent animal model in biomedical research on neuroscience, metabolic diseases, and transplant (Wernersson *et al.*, 2005; Liu *et al.*, 2014). Pig is more similar to humans in anatomy, histology, growth, development, immunity, physiology, and metabolism than rodents (Wall and Shani, 2008). A high quality draft pig genomic sequence also has become available (Walters *et al.*, 2015). Thus, pig is an ideal mammal model to investigate the mechanisms underlying fine-tuned regulation. Venø *et al.* (2015) examined the spatiotemporal regulation of circRNA in fetal porcine brain and found that introns flanking circRNAs are significantly larger than introns flanking noncircRNAs and associated with proximal complementary SINEs. Liang *et al.* (2017) identified 5 934 circRNAs in Guizhou miniature pig. They discovered 31% of circRNAs harbor well-conserved canonical miRNA seed matches, suggesting that some circRNAs act as miRNA sponges, and 149 circRNAs potentially associated with muscle growth. In the present study, we investigated the functional role of circRNAs in intermuscular fat metabolism from lean and obese pigs. We quantified the expression levels of circRNAs in lean and obese pigs by high-throughput deep sequencing.

The quality of pork consumed as food by humans has been associated the occurrence of obesity, cardiovascular diseases, diabetes, hypertension, cancer and so on. High-protein, low-fat, low-cholesterol, thin and tender, and delicious pork meat is becoming increasingly in demand among consumers. Jinhua pig is a famous local breed in China because of its good meat quality, fast fat deposition and slow growth. Landrace pig is a famous lean breed worldwide because of its fast growth, high thin meat rate, and low fat deposition (Miao *et al*. 2008; Miao *et al* 2009). These two pig species are ideal models for the comparative study of fat deposition and production differences in pigs. In the present study, we investigated the performances of Jinhua and Landrace pigs at the fattening period (at 180 days of age) and explored the expression profile of circRNAs in intermuscular fat from the two pig breeds. We prepared a comprehensive map of circRNA expression in lipid metabolism and explored the characteristics of fat deposition in mammal. A total of 5 548 circRNAs were found in the two breeds, and these circRNAs were then subjected to the Gene Ontology (GO), Cluster of Orthologous Groups of Proteins (COG), and Kyoto Encyclopedia of Genes and Genomes (KEGG) databases to explore gene regulation changes that can influence adipose tissue generation.

## 2. Results

### 2.1 Animal performances

The methods used to investigate the animal performances are presented in Supplementary Material. As shown in Table 1, the Jinhua pigs had lower initial weight, final weight, Average daily gain, average daily feed intake, dressing percentage, lean meat percentage, and *Longissimus* muscle area than the Landrace pigs (*P*< 0.05). However, the Jinhua pigs had higher feed-to-gain ratio, average backfat thickness, intramuscular fat, *a\** and *b\** than the Landrace pigs(*P*< 0.05). No significant difference in *L\**was observed between the Jinhua andLandrace pigs (*P*> 0.05). As shown in Table 1, the Jinhua pigs had poorer growth performance, but better meat quality than the Landrace pigs. In particular, the Jinhua pigs had more fat deposition at 180 days of age than the Landrace pigs.

**Table 1.** Performances in Jinhua and Landrace pigs

| Item | Number | Breeding |  | SEM | Sig |
| --- | --- | --- | --- | --- | --- |
|  |  | Jinhua pig | Landrace |  |  |
| Initial weight (kg) | 18 | 4.4 | 8.5 | 0.49 | *** |
| Final weight (kg) | 18 | 60.8 | 104.3 | 5.24 | *** |
| Average daily gain(g) | 18 | 388.77 | 643.20 | 30.70 | ** |
| Feed:gain ratio | 18 | 3.63 | 2.79 | 0.11 | ** |
| Average daily feedintake (kg) | 18 | 1.41 | 1.80 | 0.05 | ** |
| Dressing percentage(%) | 6 | 59.76 | 64.45 | 1.20 | * |
| Lean meat percentage(%) | 6 | 46.32 | 63.28 | 3.99 | ** |
| *Longissimus* muscle area (cm<sup>2</sup>) | 6 | 27.13 | 50.62 | 5.49 | ** |
| Average backfat thickness (cm) | 6 | 3.15 | 2.39 | 0.19 | ** |
| IMF (%) | 6 | 3.77 | 1.96 | 0.43 | ** |
| *L* * | 6 | 48.22 | 47.67 | 0.93 | NS |
| *a* * | 6 | 7.37 | 4.76 | 0.67 | ** |
| *b* * | 6 | 7.96 | 5.69 | 0.61 | * |

Significantly different between the two breeds, \* $P < 0.05$ , \*\* $P < 0.01$ , \*\*\* $P < 0.001$ .
*L*\* = lightness; *a*\* = redness; *b*\* = yellowness; IMF = intramuscular fat.

### 2.2 Sequencing dada and quality analysis

To construct a comprehensive map of circRNA expression in lipid metabolism in mammals, we adopted the Illumina technique for circRNA expression profile in intermuscular fat from the Jinhua and Landrace pigs at 180 days postnatal. To ensure an unbiased representation of circRNA, we refrained from doing both poly(A) selection and RNase R treatment to remove linear RNA. We depleted rRNA and then sequenced circRNAs by using the paired-end (2×100 nucleotide) Illumina technique. Data were processed to remove adapter sequences and low-quality sequences. As shown in Table 1S, 94.92 Gb clean data were obtained from the six samples of Jinhua and Landrace pigs, and the least clean data still had 13.8 Gb. Then, all reads were mapped onto the genome of *Sus scrofa* by Tophat2 software. A total of 51 912 627; 55 689 002; 47 165 141; 62 533 267; 54 782 722 and 46 305 666 reads were mapped and used for further analysis (Table S1). The GC content from the clean data was more than 53.22%. The percentage of Q30 bases was not less than 91.82%, and the data were available. As shown in Figure S1, the pink, green and blue regions in each column show the proportion of mapped reads from the exon, intergenic and intron regions in the reference genome, respectively. The height of the region represents the percentage of mapped reads in all mapped reads. The distribution of mapped reads in the Jinhua and Landrace pigs was mainly in the exon region, followed by the intergenic region. As shown in Figure S2, the circRNA junction reads in the two breeds were the most distributed on chromosome 1, followed by chromosome 15, and the least distributed on the Y chromosome, followed by chromosome 17.

### 2.3 Identification of circRNAs

CircRNAs were detected in the two breeds by using CIRI, find_circ, and CIRCexplorer. A total of 5 548 circRNAs were found, specifically 2 651 (47.78%) in the Jinhua pigs, and 2 897 (52.22%) in Landrace pigs (Table 2). The percentage of reads producing circRNAs reached 6.7%, and the percentage of reads producing circRNAs was higher in the Jinhua pigs (6.9%) than in the Landrace pigs (6.6%).

**Table 2.** CircRNA expression in Jinhua and Landrace pigs

|  | J1 | J2 | J3 | J | L1 | L2 | L3 | L | ALL |
| --- | --- | --- | --- | --- | --- | --- | --- | --- | --- |
| Number of circRNAs | 771 | 941 | 939 | 2 651 | 1 227 | 901 | 769 | 2 897 | 5 548 |
| Number of reads | 10 802 | 15 099 | 12 780 | 38 681 | 19 654 | 14 278 | 10 048 | 43 980 | 82 661 |
| Percentage of reads<br>producing circRNAs | 7.1% | 6.2% | 7.3% | 6.9% | 6.2% | 6.3% | 7.7% | 6.6% | 6.7% |

### 2.4 CircRNA differential expression analysis

Analysis showed 809 differentially expressed circRNAs between the Jinhua and Landrace pigs (Figure 1, Table 3), but only 29 of these circRNAs showed significant differences (19 upregulated and 10 downregulated). The differential expression of circRNAs between the Jinhua and Landrace pigs is shown by the volcanoe plot in Figure 1A. Each point in the volcanoe plot represents a circRNA; the greater the value, the more significant the differential expression of the circRNA is. The MA plot (Figure 1B) visually elucidates the overall distribution of the expression abundance and the multiple differences of circRNAs in the Jinhua and Landrace pigs.

**Table 3.**
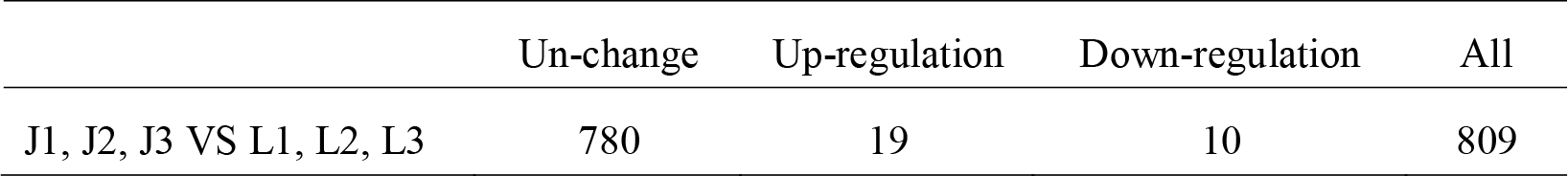
CircRNA differential expression in Jinhua and Landrace pigs

**Figure 1.**
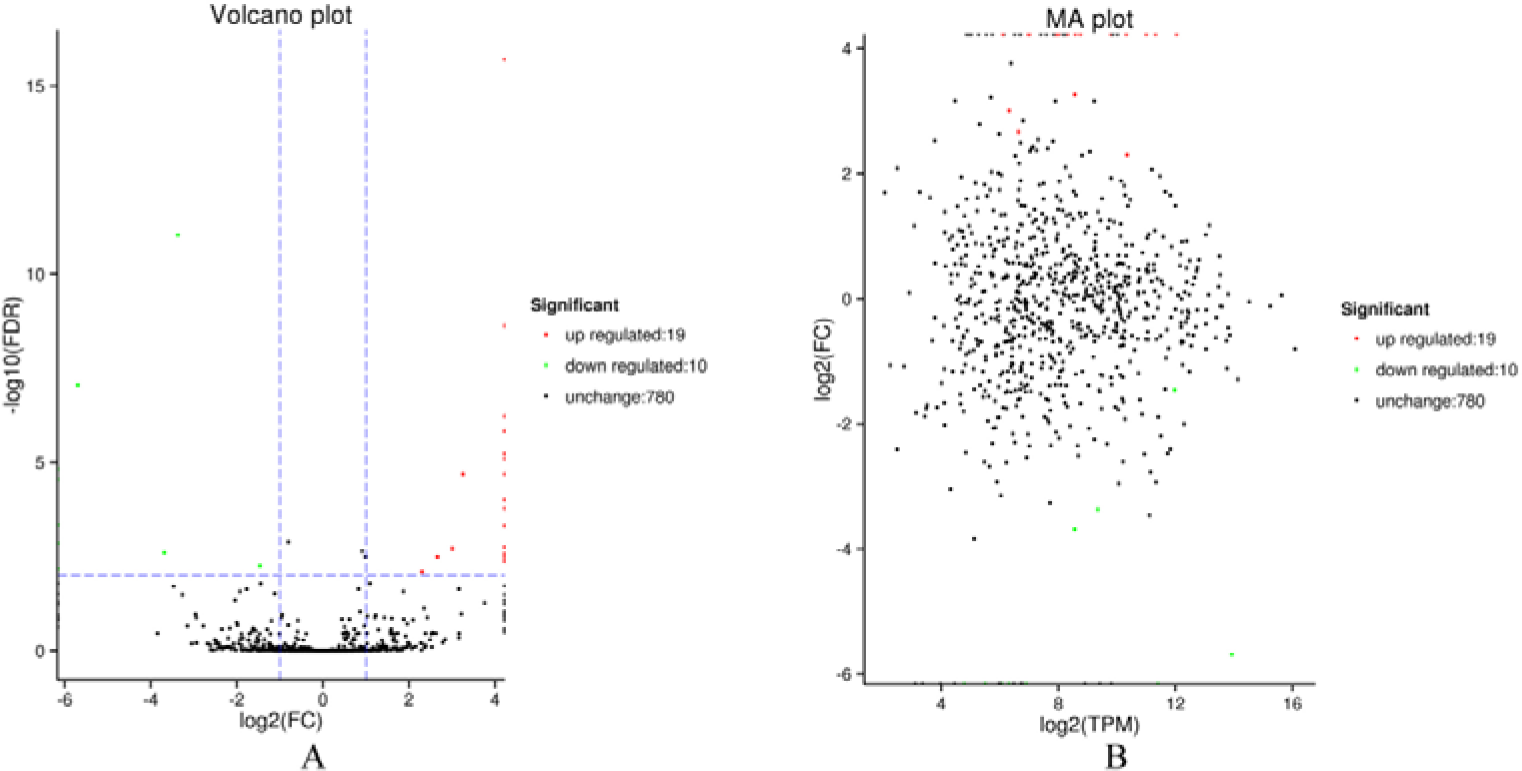
Differentially expressed circRNAs in Jinhua and Landrace pigs. Green and red dots in the plot represent circRNAs with significant differential expression. Green dots show a downregulated expression, red dots show a upregulated expression, and black dots show no significant difference. A, the abscissa indicates the logarithm of the absolute value of a circRNA in the two samples, and the differential expression of the circRNA is greater between the two samples. The ordinate indicates the negative logarithmic value of the error rate. B,the abscissa is A value, log2 (TPM), which indicates the logarithm of the mean value of circRNA expression in the two samples; the ordinate is M value, that is, log2 (FC), which indicates the logarithm of the differential times of the circRNA expression between the two breeds.

### 2.5 Clustering analysis of differentially expressed circRNAs

To quantify the expression of circRNAs in the two breeds, we clustered 5 548 circRNAs based on their expression profiles to generate a heatmap (Figure 2). Distinctive expression patterns were found between the Jinhua and Landrace pigs. All circRNAs in the Jinhua and Landrace pigs were grouped into two clusters.

**Figure 2.**
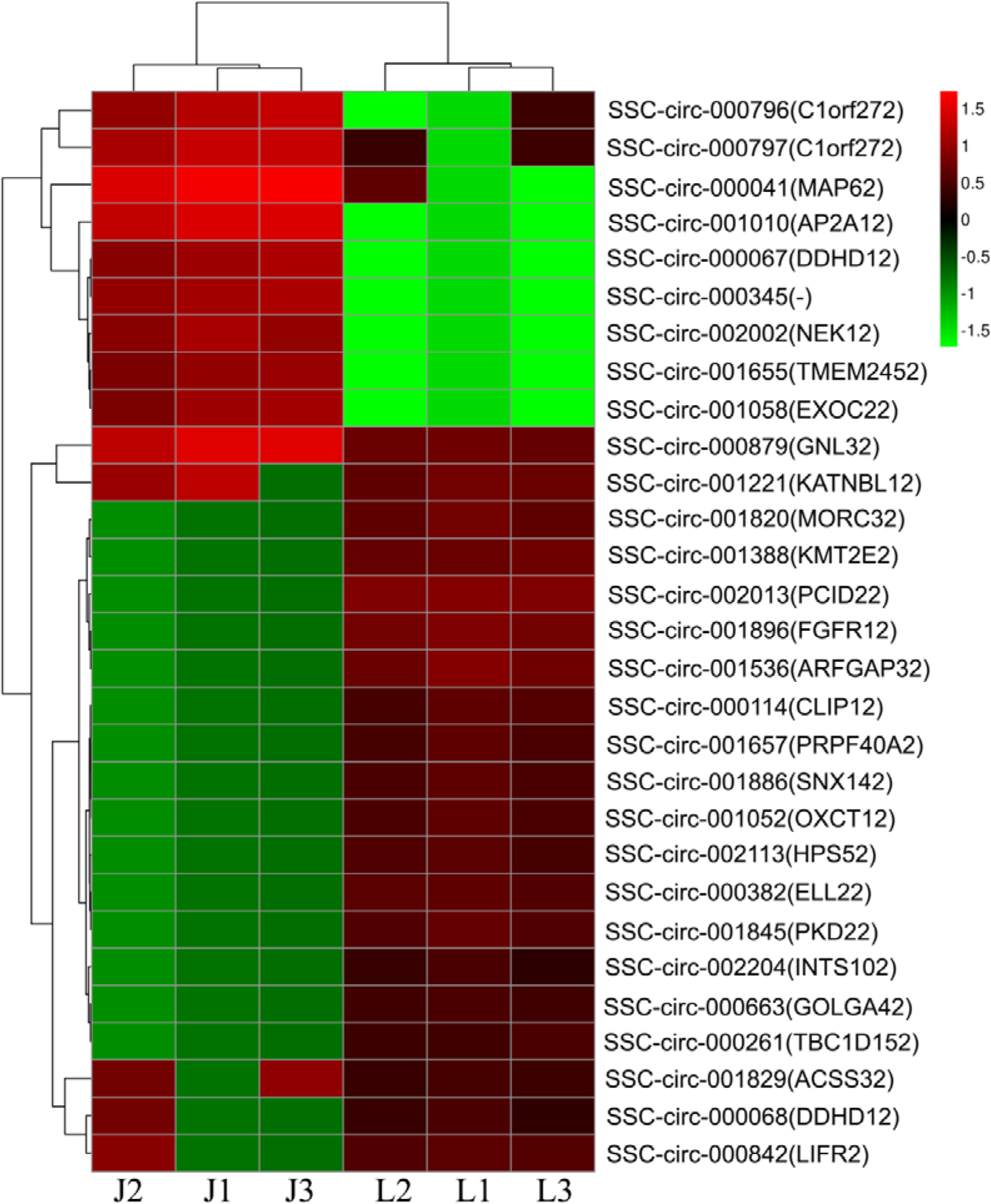
Hierarchical clustering of circRNAs in Jinhua and Landrace pigs. The columns in the chart represent different samples, and lines represent different circRNAs. Color represents the expression level of circRNA in Jinhua and Landrace pigs.

### 2.6 Functional annotation and enrichment analysis of circRNA

We analyzed the expression patterns of circRNAs in the two breeds by GO, COG, and KEGG analyses, and all the 1 376 unigenes and 27 differentially expressed genes (DEGs) were annotated (Table 4). Figure 3 shows the DEGs annotated to the GO database, and all 1306 unigenes and 27 DEGs were annotated (Table 4). Among the topGO enrichment results of circRNAs in the Jinhua and Landrace pigs (Table S2), the functions of the annotated DEGs mainly focused on protein dimerization, protein homodimerization, protein binding transcription factor, and transcription factor binding activities. On the basis of COG pathway analysis (Table 4), 544 unigenes and 12 DEGs were annotated. However, only 6%-33.33% unigenes were carried to the functional prediction (Figure 4), and they were distributed mainly to replication, recombination, and repair (3%-16.67%); lipid transport and metabolism (2%-11.11%); transcription (2%-11.11%); and signal transduction mechanisms (2%-11.11%). A total of 878 unigenes and 18 DEGs were annotated by KEGG pathway analysis (Table S3, Figure S3, S4). The genes of circRNAs were focused on the central carbon metabolism in cancer and butanoate and propanoate metabolism.

**Table 4.**
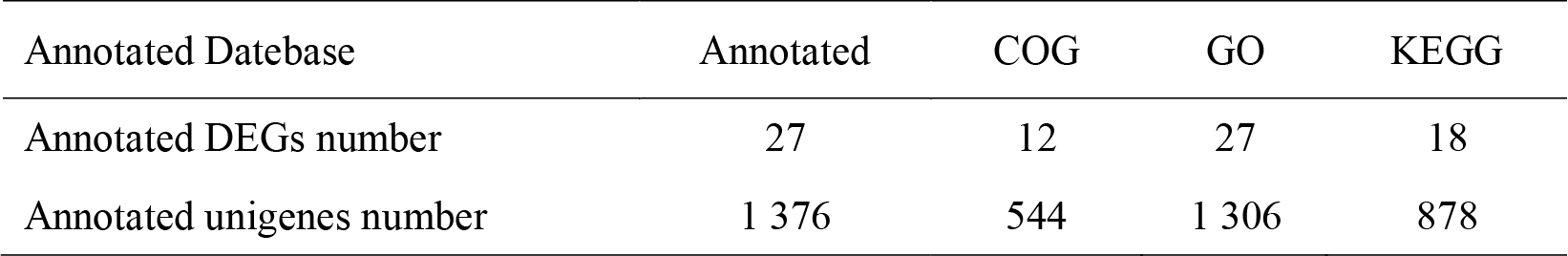
Number of new gene annotation

| Annotated Database | Annotated | COG | GO | KEGG |
| --- | --- | --- | --- | --- |
| Annotated DEGs number | 27 | 12 | 27 | 18 |
| Annotated unigenes number | 1 376 | 544 | 1 306 | 878 |

**Figure 3.**
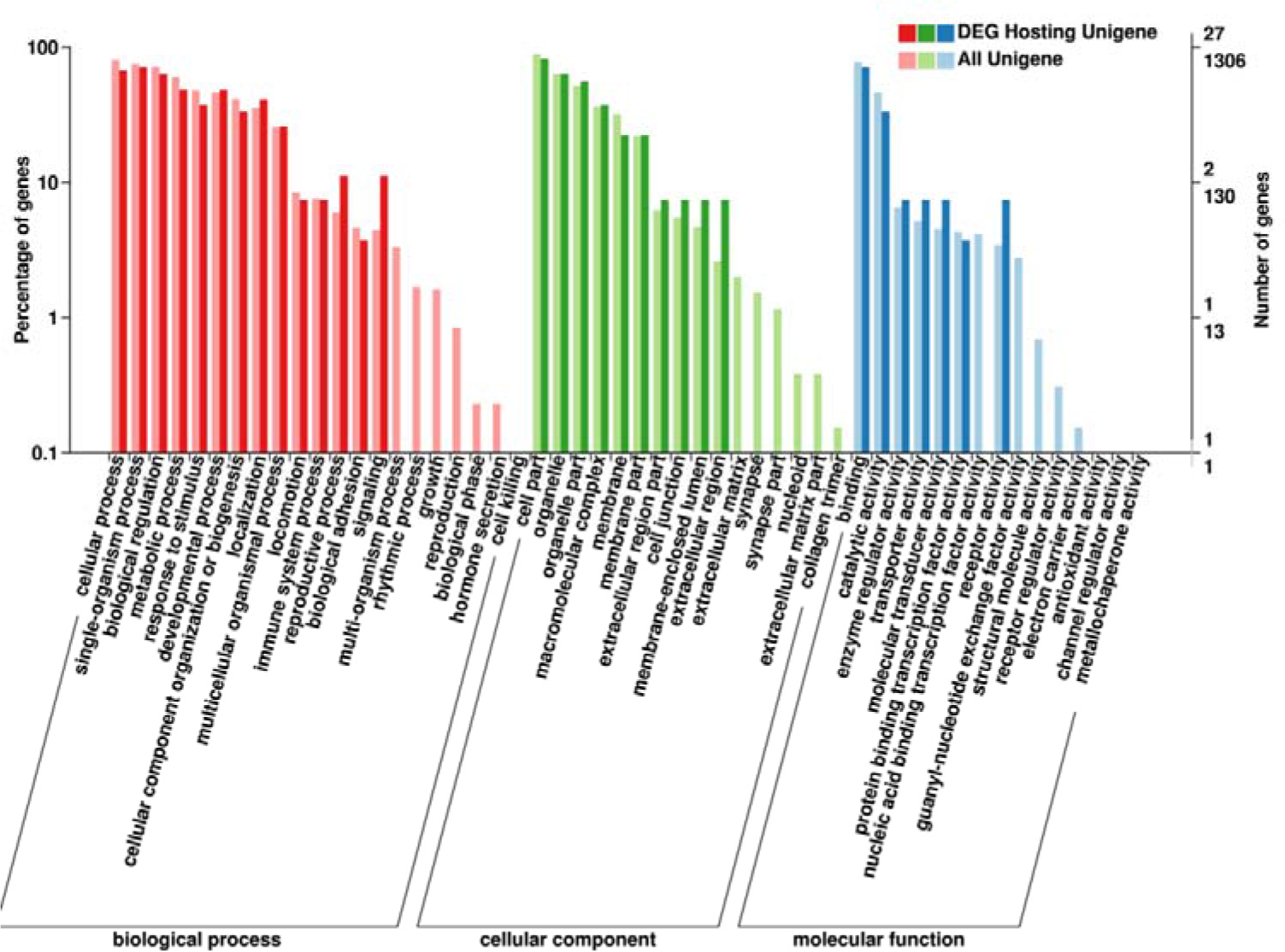
GO pathway analysis of circRNAs in Jinhua and Landrace pigs. The abscissa is classified as GO. The left side of ordinate is the percentage of the gene number, while the right side is the gene number. This image shows the functional annotation of unigenes and differentially expressed unigenes of the circRNAs, and the whole gene background.

**Figure 4.**
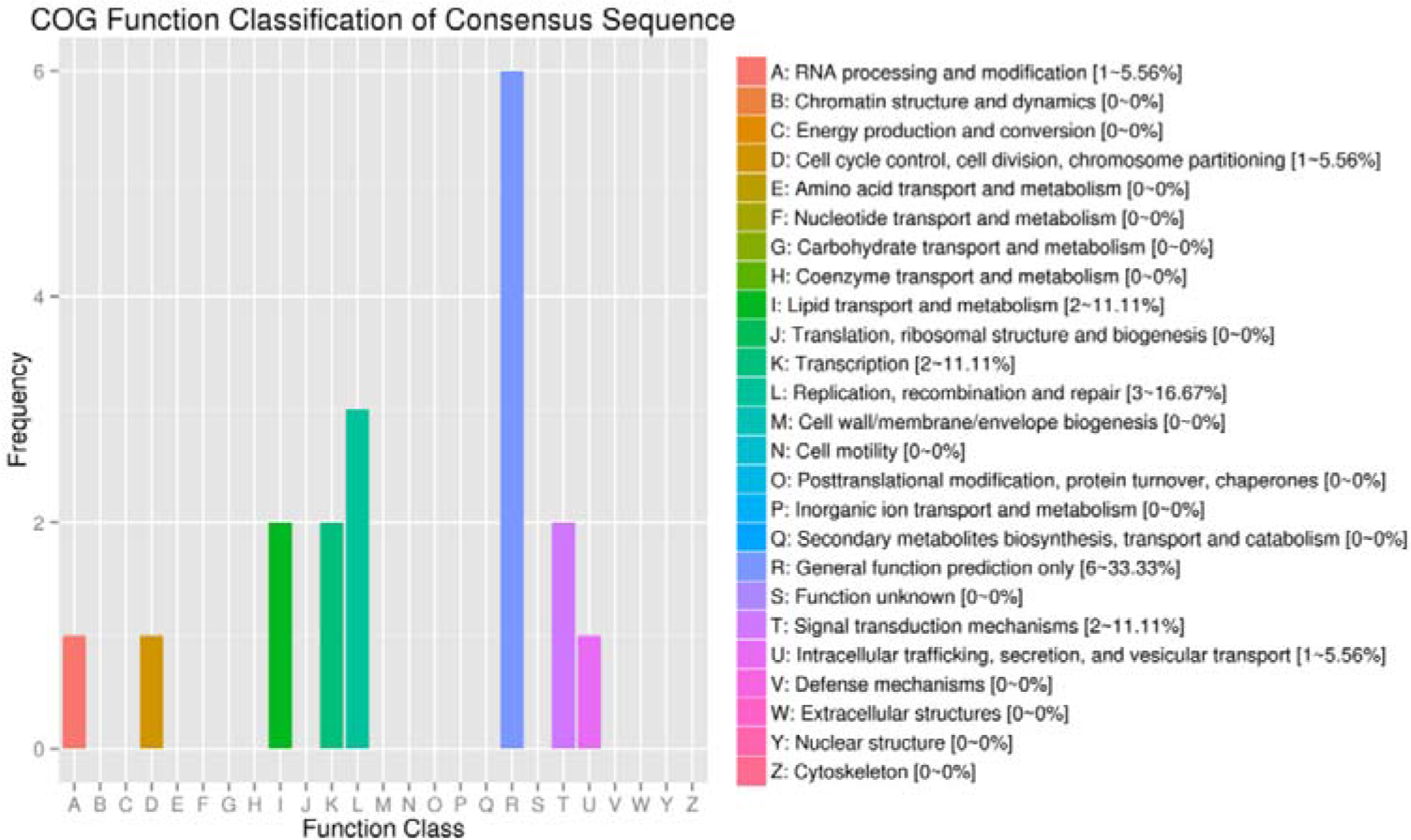
COG pathway analysis of circRNAs in Jinhua and Landrace pigs. The abscissa represents COG functional classification, and the ordinate is the number of genes. In different functional classes, the number of genes reflects the metabolic or physiological bias in the two breeds.

### 2.7 miRNA target prediction

Through RNAhybrid and miRanda software prediction, we found 550 target miRNAs with perfect seed matchs and 20 522 target genes (Supplementary miRNA data). Common *Sus scrofa* miRNAs, such as ssc-let-7 (ssc-let-7a, ssc-let-7c, ssc-let-7d, ssc-let-7e, and ssc-let-7f), ssc-miR-24, ssc-miR-103, ssc-miR-107, ssc-miR-122, ssc-miR-125, ssc-miR-127, and ssc-miR-149 were identified.

### 2.8 Analysis of expression patterns by real-time quantitative PCR

Real-time quantitative PCR results showed the differential expression patterns of 13 genes from the two breeds (Figure 5). AP2 gene was the most statistically significant in Jinhua pigs, which was 633.03 times more than that of Landrace pigs (Figure 5). KMT2E2 gene was the highestin Landrace pigs, which was 395.68 times more than that of Jinhuapigs (Figure 5).

**Figure 5.**
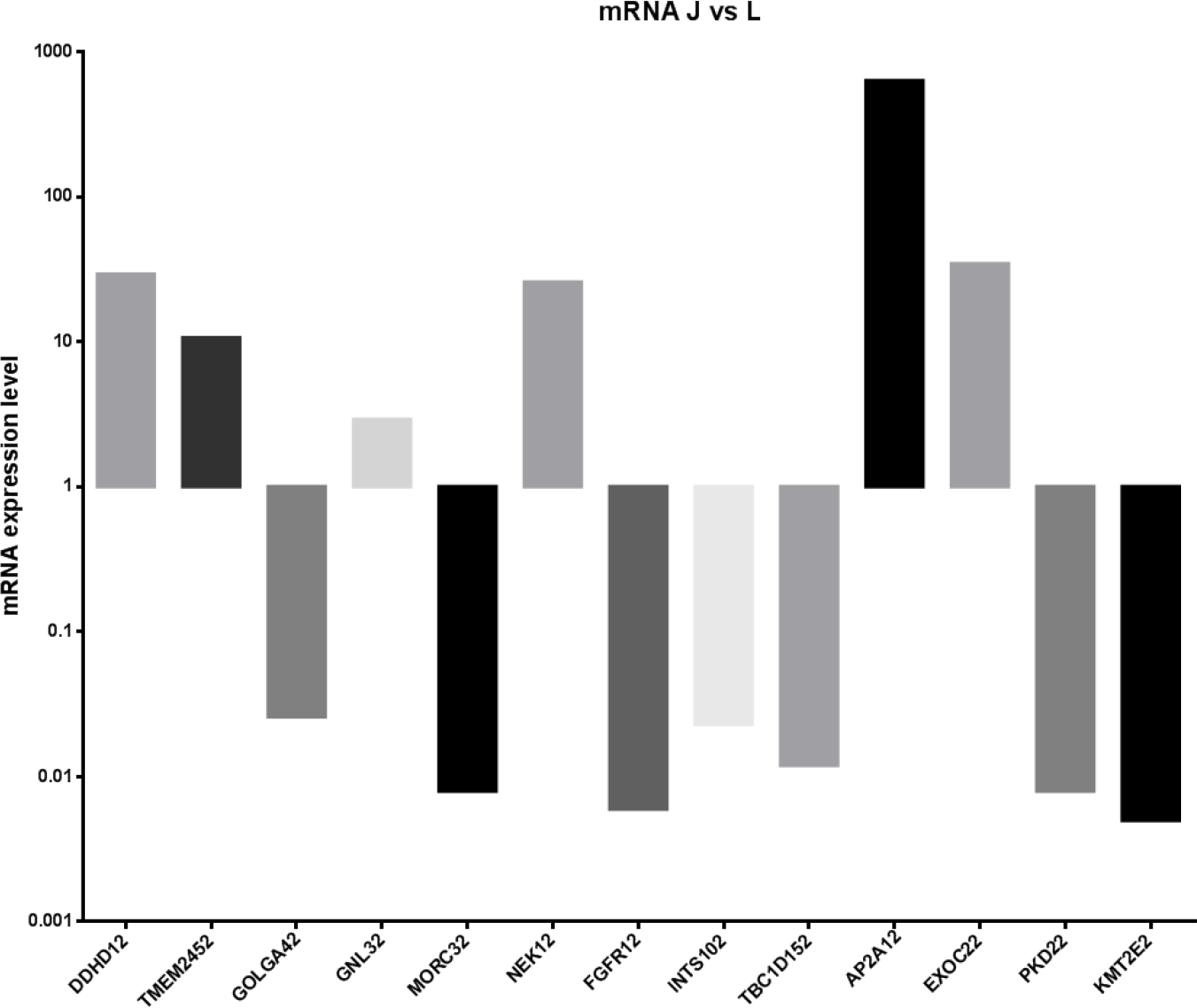
Expression levels of 13 selected genes detected by real-time quantitative PCR

## 3. Discussion

### 3.1 Animal performances

Jinhua pig is an important Chinese local breed characterized by low growth rate and high adiposity, and Landrace pig is known for its high growth rate and low fat deposition. Miao *et al*. (2008 and 2009) investigated the developmental changes of carcass composition, meat quality, and organ and lipid metabolism in Jinhua and Landrace pigs at 35–125 days of age. They found that Jinhua pigs exhibit faster fat deposition but better quality than Landrace pigs. Hormone and growth factors in lipid metabolism could induce the differences in meat characteristics in the two breeds by inhibiting fat synthesis through reducing lipogenic enzyme activities and promoting fat degradation by increasing HSL activity. We explored the performances of Jinhua and Landrace pigs at the fattening period (at 180 days of age) and found the a significant difference in fat deposition between the two breeds. Hence, the two breeds are appropriate models for investigating the mechanisms underlying the fine-tuned regulation of fat metabolism.

### 3.2 Functional analysis of circRNA

Precise gene expression in a spatiotemporal manner that is correlated with tissue-specific events is important in animal metabolism and development (Lee *et al.*, 2017; Huang and Shan, 2015). CircRNA is predominantly produced by back-splicing reactions and confirmed to serve as a transcription regulator, significantly increasing the complexity of regulation. Pig is an important domesticated farm animal that provides protein for humans. The domestic pig is increasingly being used as a nonrodent animal model in biomedical research. Venø *et al.* (2015) identified 4 634 unique circRNAs expressed from 2 195 host genes in five brain tissues across fetal porcine development, and approximately 20 % of porcine splice sites involved in circRNA production are functionally conservative between mice and humans. Liang *et al.* (2017) conducted total RNA sequencing across nine organs in Guizhou miniature pigs in three developmental stages and identified 5 934 *Sus scrofa* circRNAs. To yield a comprehensive map of circRNA expression in lipid metabolism in mammal, we investigated the circRNA expression profile in intermuscular fat from lean (Landrace) and obese (Jinhua) pigs. The study showed 5 548 circRNAs, specifically 2 651 (47.78%) in the Jinhua pigs, and 2 897 (52.22%) in the Landrace pigs. We observed 809 differentially expressed circRNAs, but only 29 of these circRNAs showed significant differences (19 upregulated and 10 downregulated). The differences revealed a spatiotemporal manner of circRNA expression, which improved the growing catalogue of annotated mammalian circRNAs. To demonstrate the function of circRNAs, we performed GO, COG and KEGG analysis on circRNA hosting genes. These analyses showed that all 1306 unigenes and 27 differentially expressed unigenes were annotated. These results indicated that the hosting genes of circRNAs are involved in lipid transport and metabolism, replication, recombination and repair, and signaling pathway. These results suggest that the functional activities of these unigenes and differentially expressed unigenes mainly correlated with *Sus scrofa* adipose metabolism and development.

### 3.3 CircRNA and miRNA interaction

CircRNA acts as a miRNA sponges (Hansen *et al.*, 2013). Hansen *et al.* (2013) showed that the circRNA sponge for miR-7 contains more than 70 conserved miRNA target sites and intensely inhibits miR-7 activity, which allows mRNAs to escape degradation following miRNA binding and then increases the levels of miR-7 targets. In the present paper, we found thatthetarget miRNA of ssc-circ-001010 is ssc-miR-24, and its target is AP2. AP2is primarily expressed in adipocytes and plays an important role in white adipocyte differentiation and lipid metabolism (Rui, 2017; Vegiopoulos, *et al.*,2017). let-7 was one of the first miRNAs discovered in *Caenorhabditis elegans* and was originally known as a regulator of developmental timing in this species (Reinhart *et al.*, 2000). Increasing studies have verified that let-7 family members are generally required to promote differentiation during development (Lee *et al.*, 2016; Caygill and Johnston, 2008; Sun *et al.*, 2016). We also found that ssc-circ-001766 and its target miRNA, ssc-let-7, possibly play important roles in adipose metabolism and differentiation. MiR-103 and miR-107 play regulatory roles in the development of adipose tissue in humans (Polster *et al.*, 2010; Bork-Jensen *et al.*, 2014). Previous studies confirmed that circRNAs interact with miRNAs to significantly regulate animal development and metabolism (Sun *et al.*, 2017; Jiang *et al.*, 2018). Our result also showed that these circRNAs and miRNAs have a complicated interaction and jointly play significant regulatory roles in *Sus scrofa* adipose metabolism and development. However, their explicit regulatory network is unclear, and insights into the regulatory network could improve understanding on animal adipose metabolism and development.

## 4. Materials and methods

### 4.1 Animals, diets and sample collection

All animals used in this study were performed according to the Regulations for the Administration of Affairs Concerning Experimental Animals (Ministry of Science and Technology, China) and approved by the Institutional Animal Care and Use Committee in Henan Institute of Science and Technology, Henan, China.

A total of 36 healthy purebred Jinhua and Landrace piglets (each breed was sex balanced and was weaned at 28 days of age) were allocated to two groups according to breed. Each group has three replicates with six piglets (sex balanced) in each replicate. The feeding experiment lasted for 145 days after 7 days of adaptation period. The pigs were reared in the same conditions, and had *ad libitum* access to basal diet and water via nipple drinkers. Experimental diets based on corn and soybean meal were formulated with CP concentrations, trace minerals and vitamins to meet or exceed the National Research Council recommendations for the different growth phases. Chemical analyses of the basal diet were carried out in accordance with the methods of AOAC (1990). From 28 days of age to 180 days of age, the corn-soybean meal diet was offered to pigs as shown in Table S4. At 180 days of age, six purebred piglets (sex balanced) of each breed were randomly selected and slaughtered by electrical stunning to determine carcass composition in accordance with the methods described by Miao *et al*. (2008 and 2009). Three pigs from Jinhua and Landrace breeds (Jinhua pig: J1, J2 and J3; Landrace: L1, L2 and L3) were randomly selected. Their intermuscular fat tissues were sampled and frozen in liquid nitrogen and then stored at −80 °C for circRNA analysis.

### 4.2 RNA extraction and detection

Total RNAs were isolated from intermuscular fat tissue by RNAprep Pure Tissue Kit following the manufacturer’s instructions (Tiangen Biotech Co., Ltd., Beijing, China). The purity, concentration, integrity and genomic DNA pollution of RNA samples were detected with a NanoPhotometer® spectrophotometer (IMPLEN, Westlake Village, CA, USA), a Qubit® RNA Assay Kit in Qubit®2.0 Fluorometer (Life Technologies, Carlsbad, CA, USA), an Aglient Bioanalyzer 2100 System (Agilent Technologies, Santa Clara, CA, USA), and electrophoretic methods to obtain high-quality samples. After assessing the quality and quantity, the total RNAs were used as templates for subsequent reverse transcription.

### 4.3 Illumina high-throughput RNA sequencing and exploration of circRNA

EpiCenter Ribo-ZeroTM Kit ((Illumina, Inc., San Diego, CA, USA) was used to remove ribosomal RNA from total RNA. After the removal of ribosomal RNA and then building a library, high-throughput RNA sequencing was performed. CircRNAs were quantitatively analyzed by Biomarker Technologies (Beijing, China). Clean reads were aligned to the reference genome (ftp://ftp.ensembl.org/pub/release-75/fasta/sus_scrofa/) by Bowtie2 (http://bowtie-bio.sourceforge.net/bowtie2/manual.shtml) (Memczak *et al*. 2013). Only mapped reads were used for analysis. Quality control of sequencing data is shown in Supplementary Material.Finally, circRNAs were verified using CIRI (Gao *et al*. 2015), Find_circ (Memczak *et al*. 2013), and CIRCexplorer (Zhang *et al*. 2014).

### 4.4 Differentially expressed genesanalysis and gene functional annotation

DEGs between the two breeds were analyzed using the method described by Audic and Claverie (1997). Differential expression analysis was performed using EBSeq software (Leng *et al.* 2013). GO enrichment analysis of DEGs was implemented using topGO software (http://www.bioconductor.org), and the statistical enrichment of DEGs in KEGG pathways was tested using KOBAS software (Mao *et al.* 2005).

### 4.5 miRNA target prediction

We used miRanda (John *et al.* 2004) and RNAhybrid (Kruger and Rehmsmeier, 2006) software to predict the miRNA sites in circRNAs with default parameters and investigate putative interactions between miRNAs and circRNAs by using only perfect seed matching without gaps of Wooble pairing.

### 4.6 Real-time quantitative PCR analysis

14genes were selected for real-time quantitative PCR analysis to confirm the results of sequencing analysis. Total RNAs were extracted using TRIzol reagent (Invitrogen, Carlsbad, CA, USA), and cDNAs weresynthetized using All-in-oneTM First-Strand cDNA Synthesis Kit(GeneCopocie, Inc., Rockville, MD, USA) in accordance with the manufacturer’s instructions. Real-time quantitative PCR was carried out as Mao *et al.*(2018). Gene primers were used in real-time quantitative PCR (Table S5).

### 4.7 Statistical analysis

Data were analyzed by One-Way ANOVA in SPSS17.0. LSD and Dunnett’s test were used to determine the significance among breed groups (Lang and Altman, 2015). All animal performance data are presented as mean ± SEM; a significance level of 0.05 was used. False discovery rate < 0.05 and log2 fold change ≥ 2 were set as thresholds for DEG screening. Landrace breed was used as the control group. The mRNA expression levels between the two breeds were calculated using the 2^−ΔΔct^ method (relative value of Jinhua and Landrace breeds).

